# High expression of angiotensin-converting enzyme-2 (ACE2) on tissue macrophages that may be targeted by virus SARS-CoV-2 in COVID-19 patients

**DOI:** 10.1101/2020.07.18.210120

**Authors:** Xiang Song, Wei Hu, Haibo Yu, Laura Zhao, Yeqian Zhao, Yong Zhao

## Abstract

Angiotensin-converting enzyme-2 (ACE2) has been recognized as the binding receptor for the severe acute respiratory syndrome coronavirus 2 (SARS-CoV-2) that infects host cells, causing the development of the new coronavirus infectious disease (COVID-19). To better understand the pathogenesis of COVID-19 and build up the host anti-viral immunity, we examined the levels of ACE2 expression on different types of immune cells including tissue macrophages. Flow cytometry demonstrated that there was little to no expression of ACE2 on most of the human peripheral blood-derived immune cells including CD4^+^ T, CD8^+^ T, activated CD4^+^ T, activated CD8^+^ T, CD4^+^CD25^+^CD127^low/−^ regulatory T cells (Tregs), Th17 cells, NKT cells, B cells, NK cells, monocytes, dendritic cells (DCs), and granulocytes. Additionally, there was no ACE2 expression (< 1%) found on platelets. Compared with interleukin-4-treated type 2 macrophages (M2), the ACE2 expression was markedly increased on the activated type 1 macrophages (M1) after the stimulation with lipopolysaccharide (LPS). Immunohistochemistry demonstrated that high expressions of ACE2 were colocalized with tissue macrophages, such as alveolar macrophages found within the lungs and Kupffer cells within livers of mice. Flow cytometry confirmed the very low level of ACE2 expression on human primary pulmonary alveolar epithelial cells. These data indicate that alveolar macrophages, as the frontline immune cells, may be directly targeted by the SARS-CoV-2 infection and therefore need to be considered for the prevention and treatment of COVID-19.

## 1. Introduction

The epidemic of a new coronavirus infectious disease (COVID-19) is wreaking havoc worldwide, caused by the severe acute respiratory syndrome coronavirus 2 (SARS-CoV-2). Currently, this virus has been globally spreading for over 7 months, with over 9 million confirmed cases and 480,000 deaths. Due to the lack of effective antiviral drugs, most patients may be treated only by addressing their symptoms, including reducing fevers and coughs. Preliminary results from the double-blind, randomized, placebo-controlled trial of intravenous remdesivir showed the reduced median recovery time (4 days) for hospitalized COVID-19 patients [1]. Based on this evidence, the United States Food and Drug Administration (FDA) has approved remdesivir under an emergency-use authorization for the treatment of adults and children with severe COVID-19. Despite this approved treatment, high mortality rates among patients have persisted. As remdesivir is an antiviral drug, the treatment is not sufficient to control COVID-19 [1]. To date, no pharmacological treatments have been shown effective for the treatment of COVID-19 [2,3]. Consequently, understanding the pathogenesis and finding an alternative treatment to improve clinical outcomes is extremely urgent as a global top priority.

SARS-CoV-2 is an RNA virus that displays high similarities, in both genomic and proteomic profiling, with SARS-CoV that first emerged in humans in 2003 after transmitting from animals in open-air markets in China [4]. The betacoronaviruses are divided into four lineages (i.e. A–D). Both SARS-CoV and SARS-CoV-2 belong to lineage B, with single strand RNA (29, 751 and 29,903 bp respectively) [4,5]. Most viruses enter cells through pattern recognition receptors (PRR) mediated endocytosis. Angiotensin-converting enzyme 2 (ACE2), with a multiplicity of physiological roles such as a negative regulator of the renin-angiotensin system [6], has been recognized as the PRR for SARS-CoV-2 infecting host cells [7], which is similar to the SARS-CoV [8]. The ACE2 expression has been mainly distributed in microvilli of the intestine and renal proximal tubules, gallbladder epithelium, testicular Sertoli cells and Leydig cells, glandular cells of seminal vesicle and cardiomyocytes [9]. The human respiratory system is primarily affected by the SARS-CoV-2 infection. Using the polyclonal anti-serum-based immunohistochemistry, the expression of ACE2 was reported on type II alveolar epithelial cells [10]. However, a single cell-RNA profiling analysis showed that only 1.4% of lung type II alveolar epithelial cells expressed ACE2 at RNA level [11], with no expression of ACE2 protein [9]. Thus, there are fundamental knowledge gaps underlying the pathogenesis of COVID-19 that need to be clarified. To understand the immunopathology and advance the strategies for the prevention and treatment of COVID-19, we examined the levels of ACE2 expression on different types of immune cells. Our data demonstrated that the activated macrophages and alveolar tissue macrophages, among others, displayed high levels of ACE2, while most of the immune cells were negative or displayed very low expressions of ACE2. This data highlights the importance of macrophages in the pathogenesis and treatment of COVID-19.

## 2. Materials and methods

### 2.1. PBMC isolation

To determine the expression of ACE2 on different types of immune cells, human buffy coat blood units (N = 6; mean age of 42 ± 13.89; age range from 27 to 58 years old; 3 males and 3 females) were purchased from the New York Blood Center (New York, NY, USA). Human buffy coats were initially added to 40 ml chemical-defined serum-free culture X-VIVO 15™ mediums (Lonza, Walkersville, MD, USA) and mixed thoroughly with 10ml pipette. Next, they were used for isolation of peripheral blood-derived mononuclear cells (PBMCs). PBMCs were harvested as previously described [12]. Briefly, mononuclear cells were isolated from buffy coats blood using Ficoll-Paque™ PLUS (γ=1.007, GE Healthcare, Chicago, IL, USA). Next, the red blood cells were removed using ACK Lysing buffer (Lonza, Walkersville, MD, USA). After three washes with saline, the whole PBMC were utilized for flow cytometry. To isolate monocytes, monocytes were purified from PBMC by using CD14^+^ microbeads (Miltenyi Biotec, Bergisch Gladbach, Germany) according to the manufacturer’s instruction, with purity of CD14^+^ cells > 90%.

### 2.2. Culture of Th17 cells

To generate Th17 cells, CD4^+^ T cells were initially isolated from PBMC using CD4^+^ microbeads (Miltenyi Biotec, Bergisch Gladbach, Germany) according to the manufacturer’s instruction, with purity of CD4^+^ T cells > 95%. The purified CD4^+^ T cells were seeded at 1 × 10^5^ cells/well in the anti-CD3 monoclonal antibody (mAb) (10 μg/mL, BD Pharmingen, Franklin Lakes, NJ, USA) precoated 96-well tissue culture-treated plate, in the presence of soluble anti-CD28 mAb (1 μg/mL, BD Pharmingen, Franklin Lakes, NJ, USA), interleukin (IL)-6 (10 ng/mL, Biolegend, San Diego, CA, USA), IL-1β (10 ng/mL, Biolegend, San Diego, CA, USA), transforming growth factor (TGF)-β1 (1 ng/mL, Biolegend, San Diego, CA, USA), IL-23 (10 ng/mL, Biolegend, San Diego, CA, USA), penicillin-streptomycin (10 μg/mL, Sigma, Saint Louis, MO, USA), the neutralizing antibodies anti-IL-4 mAb (10 μg/mL, BD Pharmingen, Franklin Lakes, NJ, USA) and anti-IFN-γ mAb (10 μg/mL, BD Pharmingen, Franklin Lakes, NJ, USA), in X-VIVO 15 serum-free medium (200 μl per well), at 37 °C, 5% CO_2_ conditions. After culturing for 5 days, the cells were stimulated with the Cell Activation Cocktail of 1.5 mM monensin sodium salt, 0.0405 mM phorbol 12-myristate 13-acetate and 0.67 mM ionomycin calcium salt (Bio-Techne Corporation, Minneapolis, MN, USA) at 2 μL. Stock solution was diluted with 1 mL of the culture medium for 4 hours at 37 °C, 5% CO_2_ conditions. Finally, the Th17 cells were harvested for flow cytometry analysis.

### 2.3. Isolation of mitochondria from platelets

Adult human platelet units (N = 3; mean age of 45.67 ± 14.98; age range from 29 to 58 years old; 2 males and 1 female) were purchased from the New York Blood Center (New York, NY, http://nybloodcenter.org/). The mitochondria were isolated from platelets using the Mitochondria Isolation kit (Thermo scientific, Rockford, USA) according to the manufacturer’s recommended protocol [13]. The concentration of mitochondria was determined by the measurement of protein concentration using a NanoDrop 2000 Spectrophotometer (ThermoFisher Scientific, Waltham, MA, USA).

For mitochondrial staining with fluorescent dyes, mitochondria were labeled with MitoTracker Deep Red FM (100 nM) (Thermo Fisher Scientific, Waltham, MA) at 37°C for 15 minutes according to the manufacturer’s recommended protocol, followed by two washes with PBS at 3000 rpm × 15 minutes [13]. Finally, the mitochondria were harvested for flow cytometry analysis.

### 2.4. Isolation of monocytes from PBMC and macrophage polarization

To generate M2 macrophages and determine ACE2 expression, monocytes were purified from human PBMC by using CD14 microbeads (Miltenyi Biotec, Bergisch Gladbach, Germany) according to the manufacturer’s instruction, with the purity of CD14^+^ cells > 90%. The isolated monocytes were seeded in the six-well tissue culture-treated plate at 5 × 10^5^ cells/ well in 2 mL X-VIVO 15 serum-free culture media at 37 °C, 5% CO_2_ condition. After 2-hour culturing, the attached monocytes were washed twice with PBS to remove floating cells and cellular debris, followed by treatment with 50 ng/mL MCSF (Sigma, Saint Louis, MO, USA) in X-VIVO 15 serum-free medium at 37 °C, 5% CO_2_. After culturing for 7 days, the M2 macrophages were treated with 1 μg/mL lipopolysaccharides (LPS) (Sigma, Saint Louis, MO, USA) or 40 ng/mL IL-4 (Biolegend, San Diego, CA, USA) in duplicates respectively for 24 hours in X-VIVO 15 serum free medium. Consequently, cells from different groups were collected for evaluations.

### 2.5. Flow cytometry

Phenotypic characterization of PBMC, monocytes, macrophages and Th17 cells were performed by flow cytometry with associated markers including: PC5.5-conjugated anti-human CD3 mAb, ECD-conjugated anti-human CD3 mAb, APC-conjugated anti-human CD4 mAb, Krome Orange-conjugated anti-CD8 mAb, APC-Alexa Fluor 750-conjugated anti-CD66b mAb, PC5.5-conjugated anti-human CD19 mAb, PC 5.5-conjugated anti-human HLA-DR mAb, Krome Orange-conjugated anti-CD14 mAb, PE-conjugated anti-human CD123 mAb, PE-conjugated anti-human CD25 mAb, PE-Cy7-conjugated anti-human CD41 mAb, and FITC-conjugated anti-human CD42a mAb purchased from Beckman Coulter (Brea, CA, USA). The antibodies FITC-conjugated anti-human Hsp60, PB-conjugated anti-human CD3, and APC-conjugated anti-IL17A mAbs were purchased from Biolegend (San Diego, CA, USA). The antibodies PE-Cy7-conjugated anti-human CD11c mAb, PE-Cy7-conjugated anti-human CD56 mAb, BV-421-conjugated anti-human CD127 mAb, PE-conjugated anti-RORγT, and BV 421-conjugated anti-RORγT mAbs were purchased from BD (Franklin Lakes, NJ, USA). PE-conjugated anti-IL17F were purchased from Invitrogen (Carlsbad, CA, USA). PE-Cy7-conjugated anti-human CD196 (CCR6) and PE-Cy7-conjugated anti-human MAVS mAbs were purchased from ThermoFisher Scientific (Waltham, MA, USA).

To determine an expression of ACE2 protein on different types of PBMC, macrophages, Th17 cells, platelets or platelet-derived mitochondria, the indirectly-labeled immunostaining with mouse anti-human ACE2 mAb (Novus Biologicals, Catalogue# NBP2-80035-100 μg, Clone #AC18F, Littleton, CO, USA) was utilized in combination with above lineage-specific fluorescence-labeled mAbs. Briefly, samples were first pre-incubated with human BD Fc Block to block non-specific binding (BD Pharmingen, Franklin Lakes, NJ, USA) for 20 minutes at room temperature, before being directly aliquoted for different antibody stainings. Cells were initially incubated with mouse anti-human ACE2 mAb at 1:50 dilution, at room temperature for 30 minutes. Next, cells were washed with PBS at 400 × g 5 minutes and then stained with FITC-conjugated AffiniPure donkey or Cy5-conjugated AffiniPure donkey anti-mouse 2nd Abs (Jackson ImmunoResearch Laboratories, West Grove, PA, USA) at 1:100 dilution for 30 minutes at room temperature. Cells only with 2^nd^ Ab staining served as control. After finishing the 2^nd^ Ab staining, cells were washed with 4 mL PBS to remove residual 2^nd^ Ab. Consequently, cells were immunostained with above lineage-specific fluorescence-labeled mAbs, as previously described [13–15]. Staining with propidium iodide (PI) (BD Biosciences, Franklin Lakes, NJ, USA) was used to exclude dead cells during the flow cytometry analysis.

For intracellular staining, cells were fixed and permeabilized according to the PerFix-nc kit (Beckman Coulter, Brea, CA, USA) manufacturer’s recommended protocol. After staining, cells were collected and analyzed using a Gallios Flow Cytometer (Beckman Coulter, Brea, CA, USA), equipped with three lasers (488 nm blue, 638 red, and 405 violet lasers) for the concurrent reading of up to 10 colors. The final data were analyzed using the Kaluza Flow Cytometry Analysis Software (Beckman Coulter, Brea, CA, USA).

### 2.6. Immunohistochemistry

Female NOD/LtJ mice (aged 3 weeks) and NOD-scid mice (aged 5–6 weeks) were purchased from Jackson Laboratories (Bar Harbor, ME, USA) and maintained under pathogen-free conditions, according to a protocol approved by the institutional Animal Care Committee (ACC). Tissue samples (e.g., small intestine, lung, liver and kidney) were fixed in 10% formaldehyde, processed, and embedded in paraffin. Tissue sections were cut at 5 mm thickness. Tissue sections from NOD/LtJ mice (3 weeks old) were utilized for immunohistochemistry including lung tissue sections (n = 4), small intestine tissue sections (n = 4), liver tissue sections (n = 4), and kidney tissue sections (n = 4). Immunostaining was performed as previously described with minor modifications [16]. To block non-specific staining, sections were incubated in a buffer containing 2.5% horse serum (Vector Laboratories, Burlingame, Ca, USA) and mouse Fc Block (BD Pharmingen, Franklin Lakes, NJ, USA) for 20 minutes at room temperature. Tissue sections were initially immunostained with ACE2 rabbit polyclonal antibody (Abcam, Cambridge, MA, USA) at 1:100 dilution for 2 hours at room temperature. Next, tissue sections were stained with Cy3-conjugated AffiniPure donkey anti-rabbit 2nd Ab (Jackson ImmunoResearch Laboratories, West Grove, PA, USA) at 1:100 dilution and combined with FITC-conjugated rat anti-mouse F4/80 mAb (eBioscience, San Diego, CA, USA) at 1:100 dilution at room temperature for 1 hour. For every experiment, only tissue sections with 2^nd^ Ab staining served as negative control. Finally, the slides were covered by using mounting medium with DAPI (Vector Laboratories, Burlingame, CA, USA) and photographed with a Nikon A1R confocal microscope on a Nikon Eclipse Ti2 inverted base, using NIS Elements Version 4.60 software.

### 2.7. Statistics

Statistical analyses were performed with GraphPad Prism 8 (version 8.0.1) software. The normality test of samples was performed by the Shapiro-Wilk test. Statistical analyses of data were performed by the two-tailed paired Student’s t-test to determine statistical significance between untreated and treated groups. The Mann-Whitney U test was utilized for non-parametric data. Values were given as mean ± SD (standard deviation). Statistical significance was defined as *P* < 0.05, with two sided.

## 3. Results

### 3.1. Little to no expression of ACE2 on most human peripheral blood-derived immune cells

To explore the direct action of SARS-CoV-2 on immune cells, we examined the ACE2 expressions on different types of immune cells from human peripheral blood (n = 6). They were characterized and gated with cell type-specific surface markers [17]: CD3 for T cells, CD3^+^CD4^+^ for CD4^+^ T cells, CD3^+^CD8^+^ for CD8^+^ T cells, CD11c^+^CD14^−^ for myeloid dendritic cells (mDC), CD14^−^CD123^+^ for plasmacytoid DC (pDC), CD14 for monocytes, CD19 for B cells, CD4^+^CD25^+^CD127^low/−^ for regulatory T cells (Tregs), CD3^+^CD56^+^ for NKT cells, CD3^−^CD56^+^ for NK cells, and CD3^−^CD66b^+^ for granulocytes (Figure 1A-C). Flow cytometry demonstrated that there were no expressions of ACE2 on most types of immune cells, or with a background level (< 5%) (Figures 1D). The percentages of ACE2^+^ cells for NK and NKT cells were only 0.36% ± 0.64% and 1.74% ± 0.5% respectively (n = 6) (Figure 1D). The activated CD4^+^HLA-DR^+^ T cells displayed only 1.79% ±1.38% of ACE2^+^ cells (n = 3). The activated CD8^+^HLA-DR^+^ was only 2.16% ± 1.16% (n = 3) (Figure 1D). The data suggests that SARS-CoV-2 virus may not directly attack blood immune cells lacking the ACE2 expression.

**Figure 1.**
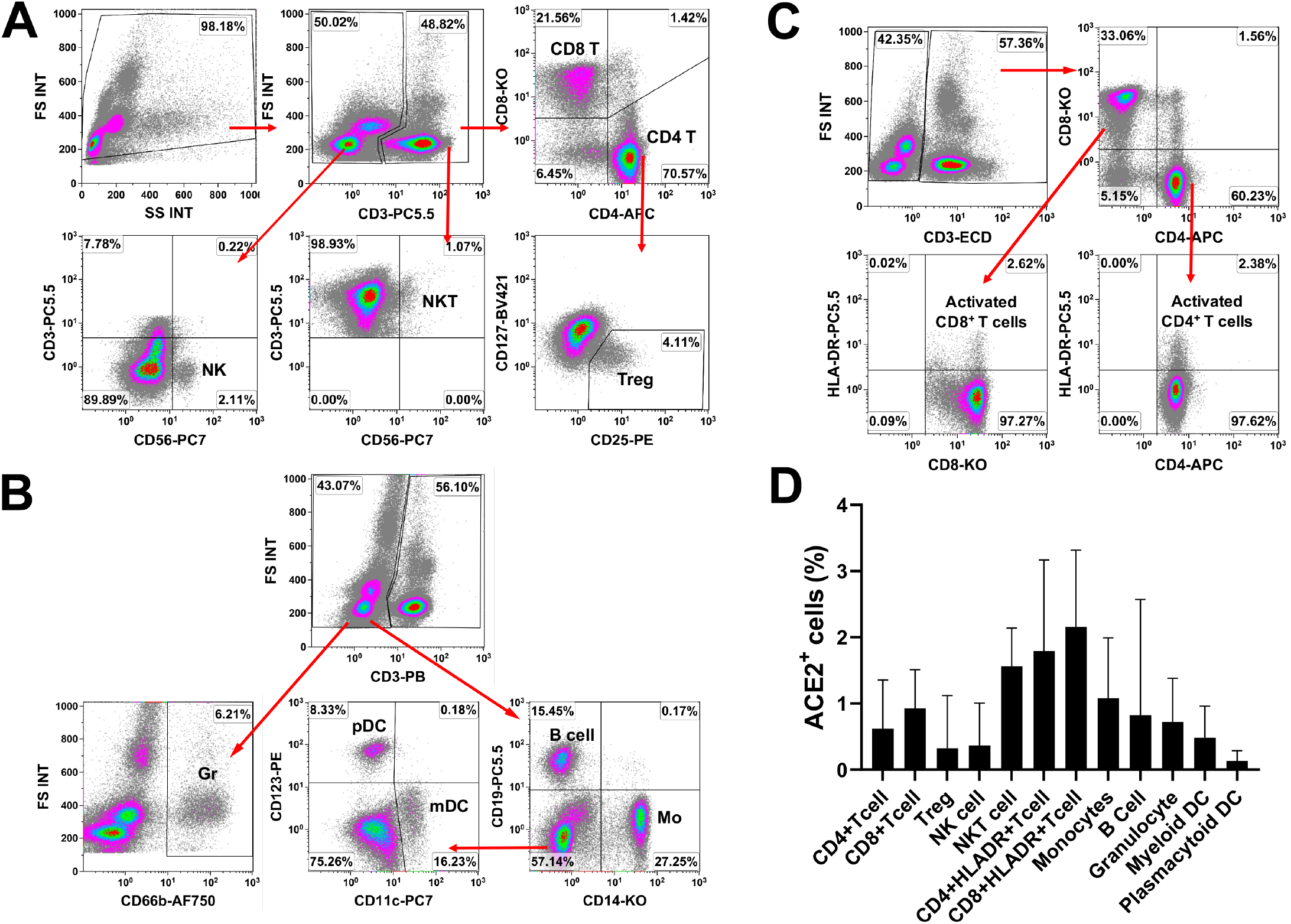
Examination of ACE2 expression on different types of immune cells. Human peripheral blood mononuclear cells (PBMC) were isolated from healthy donors (n = 6) with Ficoll-Hypaque (γ = 1.077). Red blood cells were removed using ACK lysis buffer. The remaining PBMC were utilized for flow cytometry. (**A – C**) Gating strategies of flow cytometry analysis for different types of immune cells including (**A**) CD3^+^CD4^+^ T cells, CD3^+^CD8^+^ T cells, CD4^+^CD25^+^CD127^low/−^ Treg, CD3^−^CD56^+^ NK, CD3^+^CD56^+^ NKT cells, (**B**) CD3^−^CD19^+^ B cells, CD3^−^CD14^+^ monocytes (Mo), CD14^−^CD11c^+^ myeloid dendritic cells (mDC), CD14^−^CD123^+^ plasmacytoid dendritic cells (pDC), and CD3^−^CD66b^+^ granulocytes, (**C**) activated CD4^+^HLA-DR^+^ T cells and activated CD8^+^HLA-DR^+^ T cells. (**D**) Little to no expressions of ACE2 on different gated subset of immune cells by flow cytometry. Isotype-matched IgG served as negative controls. Data are presented as mean ± SD.

### 3.2. No expression of ACE2 on Th17 cells

T-helper type 17 (Th17) cells are important pathogenic mediators for several autoimmune diseases, potentially contributing to the pathogenesis of COVID-19. RORγt (retinoic acid receptor-related orphan nuclear receptor gamma t) belong to nuclear hormone receptors (NHRs) and act as a crucial transcription factor for the differentiation and function of Th17 cells both in humans and mice [18]. Using RORγt, interleukin-17A (IL-17A), IL-17F, and CCR6 as specific Th17 markers [18], the purity of IL17A^+^RORγT^+^ Th17 cells was 19.91% ± 2.75% (Figure 2A). The percentage of IL17A^+^IL17F^+^ Th17 cells was 20.19% ± 2.94% (Figure 2A). The purity of IL17A^+^CCR6^+^ Th17 cells was 20.14% ± 2.86% (Figure 2A). The gated Th17 cells failed to express ACE2 (Figures 2B, n =4). This data implies no direct interaction between Th17 cells and SARS-CoV-2.

**Figure 2.**
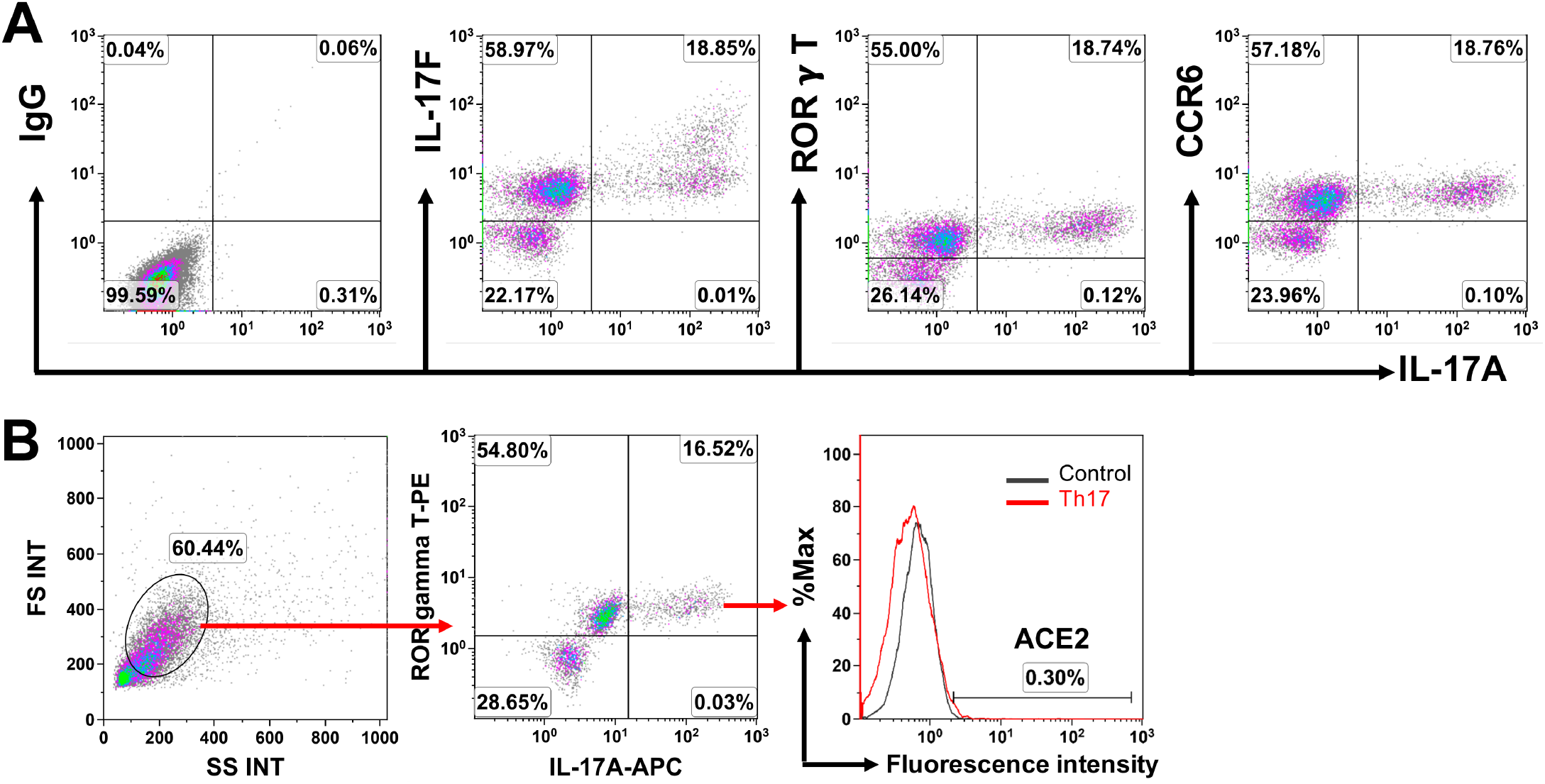
No expression of ACE2 on Th17 cells. Generation of Th17 cells was performed according to the protocol[55] with some modifications. In brief, the purified CD4^+^ T cells were seeded at 1.0–2.0 x 10^5^ cells/mL on the anti-CD3 mAb-precoated plate in the presence of anti-CD28 antibody (1 μg/mL), IL-6 (25 ng/mL), IL-1β (10 ng/mL), TGF-β1 (2.5 ng/mL), IL-23 (10 ng/mL), anti-IL-4 (10 μg/mL) and anti-IFN-γ (10 μg/mL) mAbs, in X-VIVO 15 serum-free culture media. After culture at 37℃ 5% CO_2_ conditions for 5 days, Th17 cells were utilized for experiments. (**A**) Determine the purity of Th17 cells by using Th17 cell-associated markers. Isotype-matched IgG served as negative controls. (**B**) Flow cytometry shows the level of ACE2 expression on the gated RORγT^+^IL-17A^+^ Th17 cells. Overlay of histograms show no ACE2 expression on the gated RORγT^+^IL-17A^+^ Th17 cells (red) relative to the IgG control (grey).

### 3.3. No expression of ACE2 on platelets

Increasing clinical evidence demonstrated the coagulation abnormalities in COVID-19 subjects including disseminated intravascular coagulopathy (DIC) and low levels of platelet count [19,20]. To determine whether platelets were directly targeted by SARS-CoV-2 or trigged by viral inflammatory reactions, we examined the ACE2 expression on the highly-purified CD41b^+^CD42a^+^ platelets from human peripheral blood (Figure 3A, n = 3). Flow cytometry established that there was no ACE2 expression on platelets (Figures 3B and C, n = 3), highlighting that the coagulopathy may be indirectly caused by the viral inflammation.

**Figure 3.**
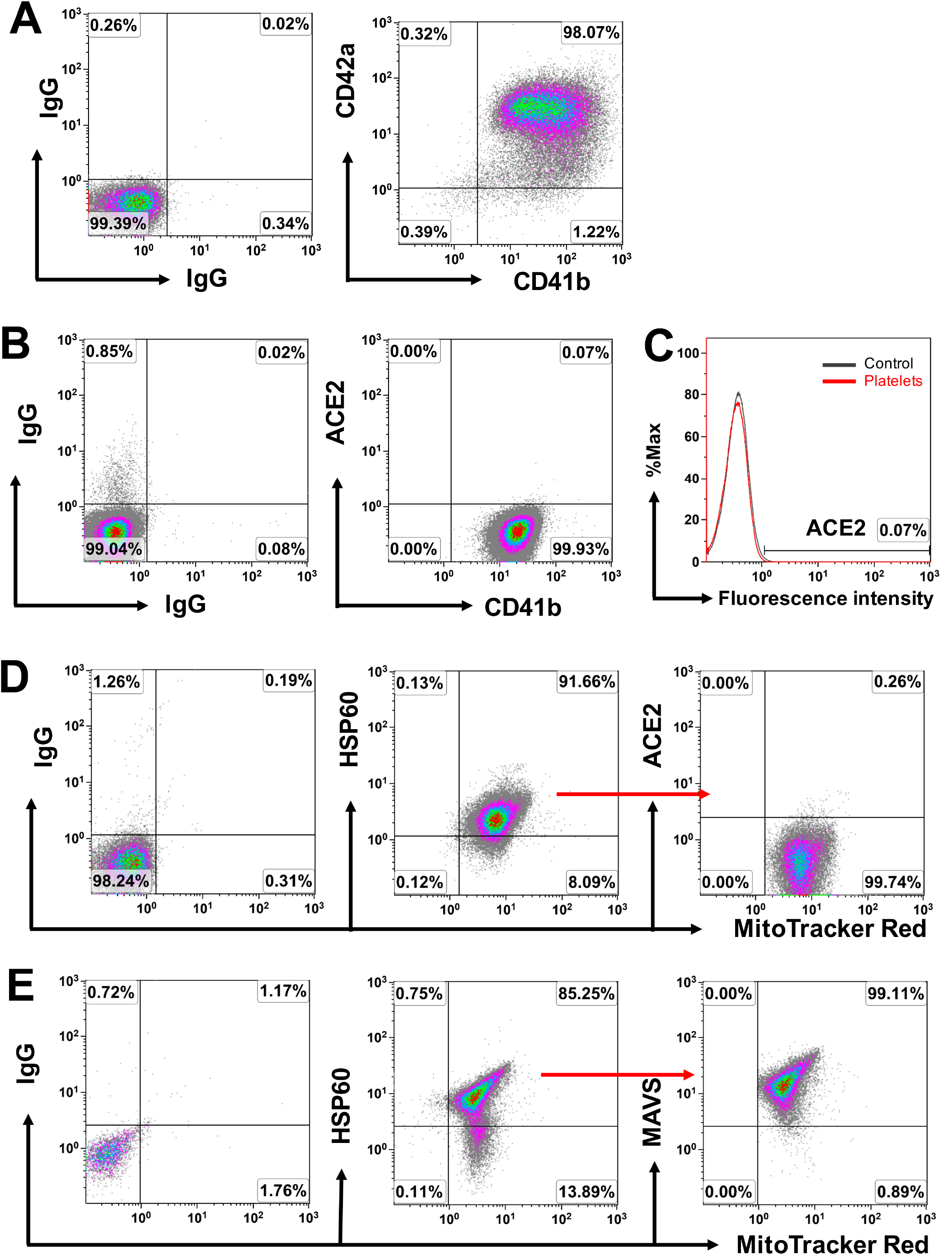
No expression of ACE2 on platelets and mitochondria. (**A**) Flow cytometry showed the expression of platelet-specific markers CD41b and CD42a on apheresis platelets, with a high purity. (**B**) Flow cytometry shows no ACE2 expression on platelets. (**C**) Overlay of histograms show no ACE2 expression on the platelets (red) relative to the IgG control (grey). (**D**) The purified platelet-derived mitochondria were labeled with MitoTracker Red and positive with mitochondrial marker HSP60, but failed to display ACE2 expression. (**E**) High expression of MAVS on platelet-derived mitochondria (n = 3). Isotype-matched IgG served as negative controls.

Our previous work established that platelets could release mitochondria contributing to the immune modulation and islet β-cell regeneration [13]. To explore the mitochondrial function in viral infection, flow cytometry indicated that while the purified platelet-derived mitochondria did not express ACE2 (Figures 3D), they strongly display the mitochondrial antiviral-signaling protein (MAVS) with the percentage of MitoTracker Red^+^HSP60^+^MAVS^+^ mitochondria at 96.02% ± 2.74% (Figure 3E, n = 3) [21]. This data suggests that platelets may have potential to improve antiviral immunity through the releasing mitochondria.

### 3.4. High expression of ACE2 on the activated type 1 macrophages (M1)

Macrophages have been characterized with type 1 macrophages (M1, inflammatory) and type 2 macrophages (M2, anti-inflammatory), according to their phenotypic differences such as spindle-like morphology and high expression CD206 and CD209 on M2 macrophages [15]. Initially, flow cytometry established that CD14^+^ monocytes from human peripheral blood failed to express ACE2 (1.78% ± 1.94%, n = 6) (Figure 4A). M2 macrophages were then generated in the presence of 50 ng/mL macrophage colony-stimulating factor (M-CSF) with the percentage of spindle-like cells at 63.25% ± 8.85% (n = 4). To evaluate the ACE2 expressions on macrophages, M2 macrophages were activated by the treatment with lipopolysaccharide (LPS) [12] and interleukin-4 (IL-4) respectively [22]. Phase contrast image showed significant differences in the morphology between two groups (Figure 4B). LPS-treated M2 macrophages exhibited pseudopod-like protrusions compared to the spindle form of IL-4-treated M2 macrophages (Figure 4B, left). Flow cytometry demonstrated that the level of ACE2 expression was higher on the LPS-activated M1 macrophages than that of IL-4-treated M2 macrophages (Figure 4C-E). This finding was further confirmed by the confocal microscopy and image analysis (Figure 4F). Therefore, the data suggests the upregulation of ACE2 expression on the activated M1 macrophages.

**Figure 4.**
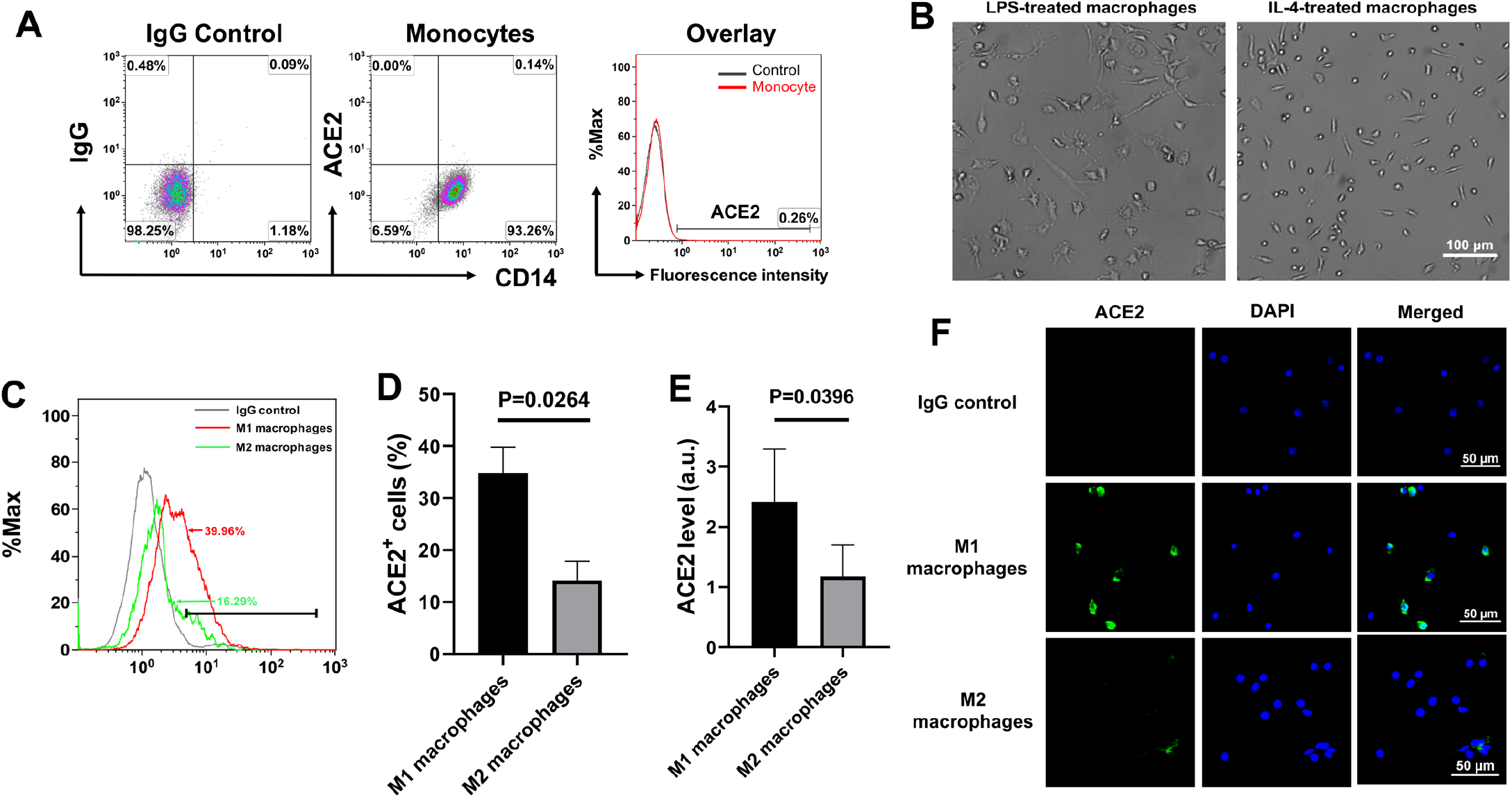
Different levels of ACE2 expressions on M1 and M2 macrophages. (**A**) The freshly-purified CD14^+^ monocytes from human peripheral blood failed to express ACE2. Isotype-matched IgG served as control. (**B – F**) The purified CD14^+^ monocytes were initially seeded in the tissue culture-treated 6-well plate at 5 × 10^5^ cells/well and cultured in X-VIVO 15 serum-free media with 50 ng/mL M-CSF[12], at 37 °C, 5% CO_2_ conditions. After 7 days, M2 macrophages were treated with 1 μg/mL LPS or 40 ng/mL IL-4 for 24 hours respectively. (**B**) Morphological change of M2 macrophages after the treatment LPS (N = 4). IL-4-treated macrophages served as control with maintaining the morphology of spindle-like cells (right). (**C**) Overlay histogram shows the high expression of ACE2 on M1 macrophages (red) in comparison with M2 macrophages (green). The 2^nd^ Ab staining served as negative control (grey). (**D**) M1 macrophages display higher percentage of ACE2^+^ cells than that of M2 macrophages. Data are presented as mean ± SD. N = 3. (**E**) M1 macrophages display higher level of ACE2 fluorescence intensity than that of M2 macrophages. Data are presented as mean ± SD. N = 3. (**F**) Confocal microscopy shows high expression of ACE2 on M1 macrophages. Representative images were from one of immunostaining with four tissue sections.

### 3.5. Expression of ACE2 on tissue macrophages

To determine the expression of ACE2 on tissue macrophages, we initially performed double-staining with mouse macrophage marker F4/80 through the immunohistochemistry in the small intestine, lung, liver, and kidney tissues of 3-week non-obese diabetic (NOD) mice. Using an expression of ACE2 in the small intestine as a positive control (Figure 5A), the data revealed that most ACE2 expressions were co-localized with the F4/80^+^ tissue macrophages in the lung and liver (Figure 5B and C), which are known as dust cells (alveolar macrophages, Figure 5B) and Kupffer cells (Figure 5C) respectively. Unexpectedly, there was little to no expression of ACE2 on the alveolar epithelial cells (Figure 5B). Additionally, we observed the strong expressions of ACE2 on the proximal tubules, with some scattered F4/80^+^ macrophages double-positive with ACE2 staining (Figure 5D).

**Figure 5.**
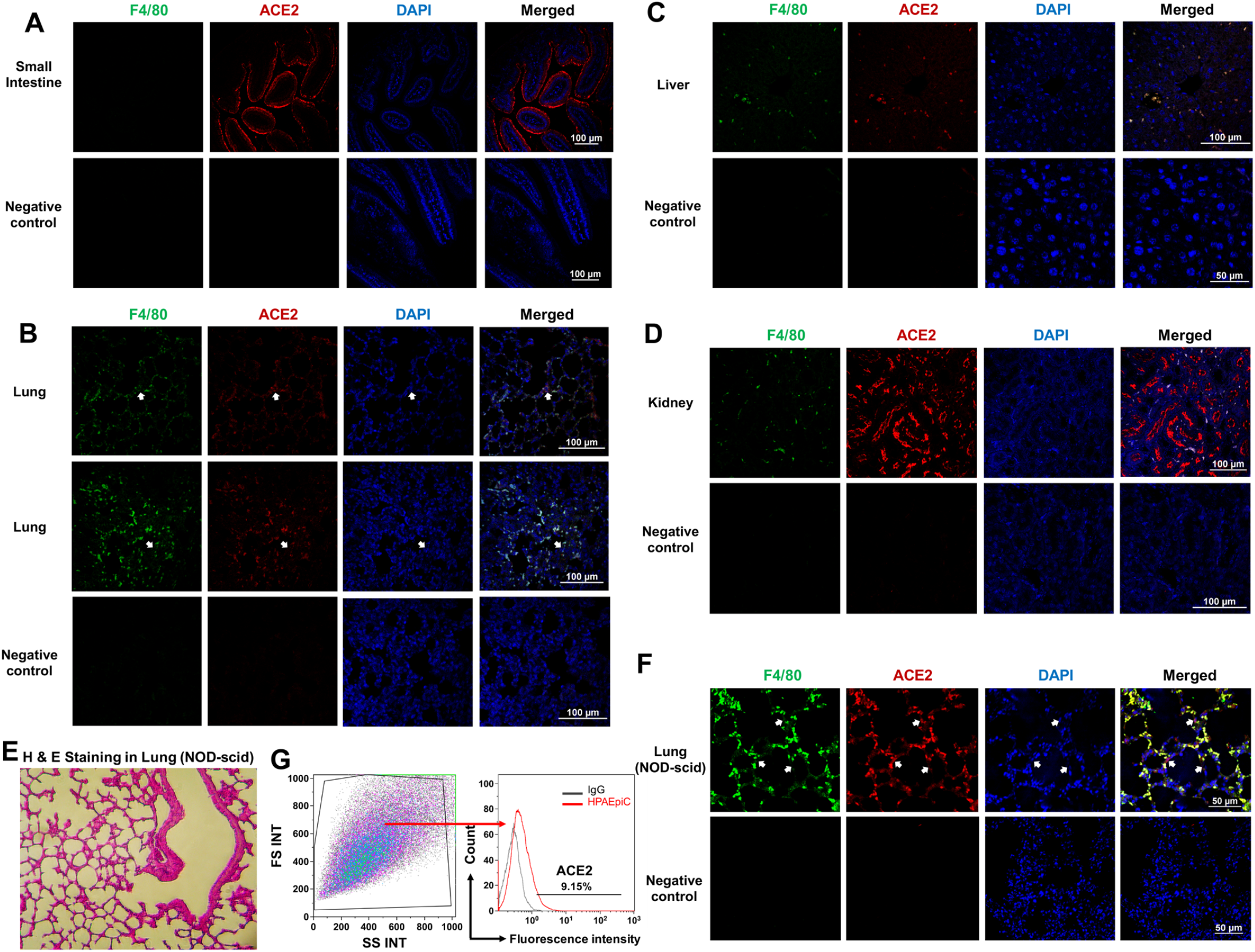
Expression of ACE2 on tissue macrophages. **(A-D)** Tissue sections were prepared from tissue samples of NOD mice (n = 6) at 3 weeks old. (**A**) High expression of ACE2 in the microvilli of the small intestine serve as positive control. (**B**) Expression of ACE2 was co-localized with F4/80^+^ alveolar macrophages of the lung. There was little to no expression of ACE2 on alveolar epithelial cells (pointed by white arrows). (**C**) Expression of ACE2 was co-localized with F4/80^+^ Kupffer cells of the liver. (**D**) High expression of ACE2 in the renal proximal tubules of the kidney, with co-localization of some ACE2^+^ cells with F4/80^+^ macrophages. (**E**) Hematoxylin and Eosin (H&E) staining of lung tissue of NOD-scid mice. Original magnification, ×200. (**F**) Expression of ACE2 was co-localized with pulmonary F4/80^+^ alveolar macrophages of NOD-scid mice (n = 4). There was low expression of ACE2 on alveolar epithelial cells (pointed by white arrows). (**G**) Analysis of ACE2 expression on the primary human pulmonary alveolar epithelial cells (HPAEpiC) by flow cytometry. The 2^nd^ Ab staining served as negative control (grey) for ACE2 immunostaining (red). The total cell population (left panel) was gated for flow cytometry analysis.

To further determine the distribution of ACE2 expression on alveolar macrophages, tissue sections were examined by utilizing non-inflammatory lung tissue from NOD-scid mice (Figure 5E). Immunohistochemistry confirmed the marked co-localization of ACE2 expression on F4/80^+^ alveolar macrophages (Figure 5F), with a low expression of ACE2 on alveolar epithelial cells. Next, we utilized the primary human pulmonary alveolar epithelial cells (HPAEpiC) to define their level of ACE2 expression. Flow cytometry proved the low level (9%) of ACE2 expression on human pulmonary alveolar epithelial cells (Figure 5G). Therefore, these data indicate the high expression of ACE2 on tissue macrophages.

## 4. Discussion

The human pulmonary system is primarily targeted by SARS-CoV-2. ACE2 and proteases such as TMPRSS2 (transmembrane protease, serine 2) or cathepsin B/L were utilized for host cellular entry of SARS-CoV-2[23]. ACE2 may act as a limiting factor for viral infection at the initial stage [24]. Sungnak et al. reported the high expression of ACE2 in nasal epithelial cells [24]. Our current studies demonstrated little to no expression of ACE2 on both primary human pulmonary alveolar epithelial cells and mouse lung alveolar epithelial cells, which is consistent with previous reports [9,24]. Notably, we found that high expression of ACE2 was colocalized with tissue macrophages of the lung (alveolar macrophages) and liver (Kupffer cells), and up-regulated on the activated M1 macrophages. However, most immune cells in human peripheral blood were negative for ACE2 expression, including the freshly-isolated monocytes. Therefore, these data highlight the importance of alveolar macrophages during the pathogenesis of lung damage in COVID-19 subjects. Based on this evidence, we propose that lung macrophages may be directly targeted by the SARS-CoV-2 and play a critical role in the initiation and development of COVID-19 (Figure 6). This perspective may advance the understanding of the clinical course of COVID-19 and facilitate the development of prevention and treatment strategies. Post viral infection, the SARS-CoV-2 may either (1) be directly cleared by the healthy macrophages with asymptomatic or mild clinical symptoms or (2) destroy the dysfunctional macrophages and evoke the immune system with cytokine storm, leading to severe clinical symptoms such as high fever, hypoxia and acute respiratory distress syndrome (ARDS) (Figure 6). To this respect, it is critical to protect and restore the functions of alveolar macrophages (or other tissue macrophages) for the prevention and treatment of COVID-19.

**Figure 6.**
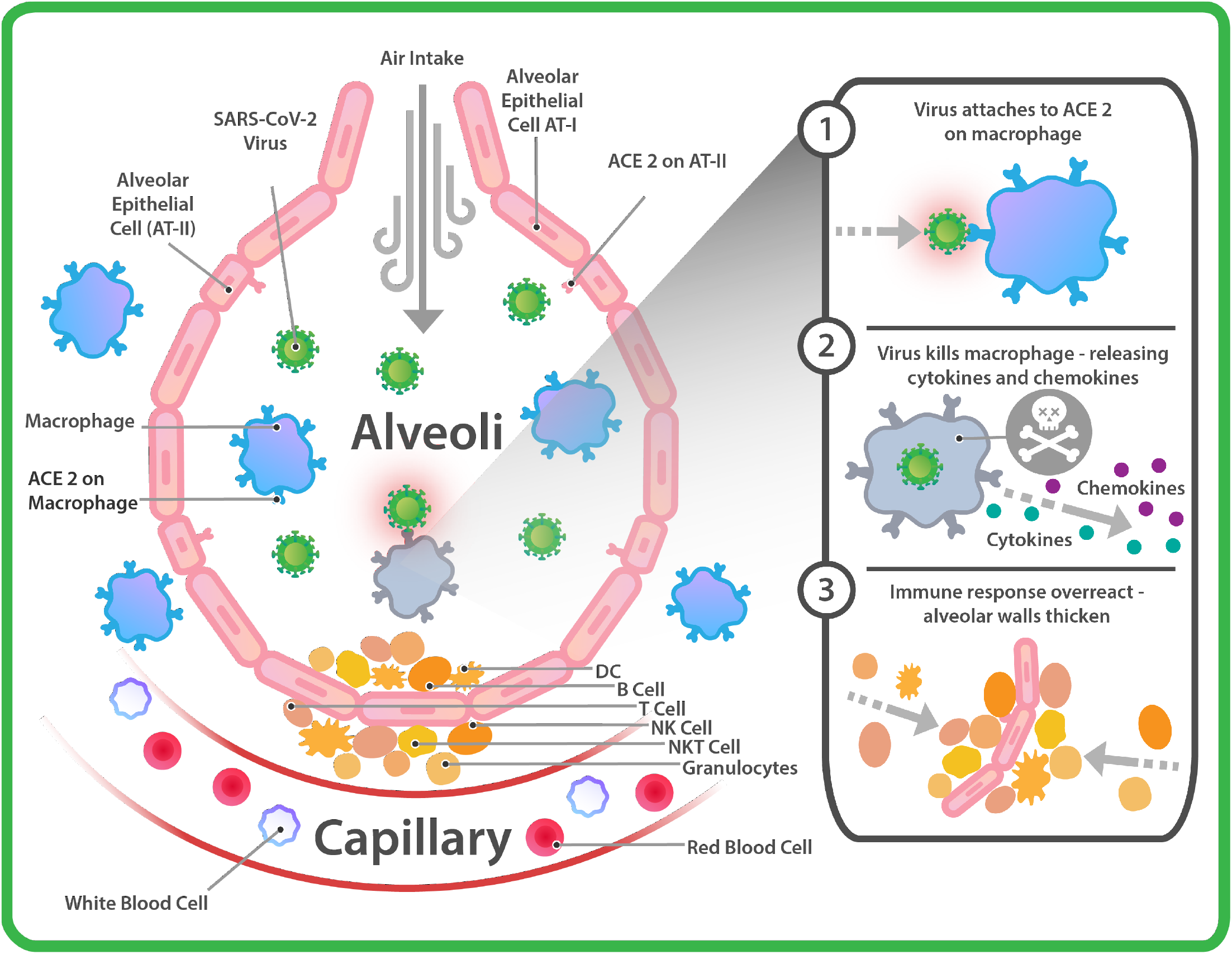
Outline the proposed mechanism underlying the pathogenesis of COVID-19. ACE2 protein was primarily displayed on alveolar macrophages of lung, with no or low expression on alveolar type 1 (AT-I) and type 2 (AT-II) epithelial cells. Upon entering the pulmonary alveoli, healthy alveolar macrophages may directly kill the virus, with asymptomatic or mild clinical symptoms. At this earlier stage 1, the infected alveolar macrophages may alternatively recruit other immune cells to build up the antiviral immunity through releasing cytokines (e.g., IL-1, IL-6, IL-12, and TNFα) and chemokines (e.g. CXCL1 and CXCL2 to recruit granulocytes, CXCL10 to recruit T cells, NK cells, and DCs). For instance, the recruited CD4^+^ T cells may secret interferon (IFN)-γto strengthen the antiviral immunity of alveolar macrophages and minimize the viral load. However, if this first line of defense is broken, the more cytokines and chemokines are released from the dead cells or dead-cell engulfed macrophages (Stages 2 and 3), the more immune cells are infiltrated into pulmonary systems, leading to patients experiencing a rapid deterioration and the development of ARDS with high fatality in the clinic.

Clinical autopsies from SARS-CoV-infected patients demonstrated that there were major pathological changes in the lungs, immune organs, and small systemic blood vessels with vasculitis. The detection of SARS-CoV was primarily found in the lung and trachea/bronchus, but was undetectable in the spleen, lymph nodes, bone marrow, heart and aorta [25]. This evidence highlights the overreaction of immune responses induced by viral infection which resulted in significant harm, as evidenced by pathogenesis of the lungs, immune organs, and small systemic blood vessels. The pathological study in COVID-19 patients revealed that the majority of infiltrated immune cells in alveoli were macrophages and monocytes, with minimal lymphocytes, eosinophils and neutrophils[26]. There were abundant proinflammatory macrophages in the bronchoalveolar lavage fluid of severe COVID-19 patients [27]. Thus, the virus-infected alveolar macrophages play a critical role in the pathogenesis of COVID-19 and SARS [28–30] and may recruit the lung infiltration of additional immune cells through predominantly releasing cytokines and chemokines [31,32], resulting in pulmonary edema and hypoxemia: the hallmark of acute respiratory distress syndrome (ARDS) (Figure 6). Consequently, the viral inflammation disrupts the homeostasis and causes multiple dysfunctions including lymphopenia [33], coagulopathy [19,34], diarrhea [33], liver injury [35], manifestations of cardiovascular [36,37] and central nervous system [38], and renal failures [33]. Additionally, the percentage of inflammation-associated Th17 cells was markedly increased in the peripheral blood of Covid-19 patients [39]. Th17 cells act as important pathogenic mediators for several autoimmune diseases including type 1 diabetes (T1D), multiple sclerosis, rheumatoid arthritis, alopecia areata (AA), psoriasis, and even the insulin resistance in type 2 diabetes (T2D) [40–42]. Thus, the viral inflammation caused by SARS-CoV-2 is very similar to the majority of autoimmune diseases in humans caused by overactive immune systems. Therefore, immune modulation strategy may be potentially beneficial to enhance anti-viral immunity and efficiently reduce the viral load, improve clinical outcomes, expedite patient recovery, and decline the rate of mortality in patients after being infected with SARS-CoV-2.

Macrophages are widely distributed in human tissues and organs with pleiotropic functions in maintaining homeostasis. Therefore, dysfunctions of macrophages may increase vulnerability to the viral infection. For example, the phenotype of intestinal macrophages may be changed during the chronic intestinal inflammation and infection [43]. Due to high expression of ACE2 in the epithelial cells of the intestine and the utilization of immune-suppression regimens, patients with chronic inflammatory bowel disease (IBD) might have an increased risk of the SARS-CoV-2 infection [44]. Additionally, increasing clinical evidence demonstrates that the chronic metabolic stress-induced inflammation causes multiple dysfunctions of macrophages, leading to the insulin resistance in type 2 diabetes [45]. Clinical studies have shown that type 2 diabetes is one of the major risk factors for COVID-19 [33,46–48]. Given this comorbidity, COVID-19 patients’ alveolar macrophages may not be efficient in fighting off the SARS-CoV-2 infection.

Alveoli are the terminal ends of respiratory tracts and function as the basic respiratory units comprising three primary cells: alveolar type 1 cells (AT-I) for building up the alveolar wall, alveolar type 2 cells (AT-II) secreting a lipoprotein surfactant that reduces the surface tension and prevents the adhesion of the alveolar walls, and alveolar macrophages. Alveolar macrophages reside in the pulmonary alveoli and the inter-alveolar septum, close to the pneumocytes (alveolar epithelial cells). The current study demonstrated little to no expression of ACE2 on pneumocytes (including AT-I and AT-II pneumocytes). Notably, peripheral blood-derived CD14^+^ monocytes did not express ACE2, as demonstrated by flow cytometry. In contrast, there are high expressions of ACE2 on alveolar macrophages, implying that the circulating monocytes may differentiate into alveolar macrophages after migration into the pulmonary tissues with an up-regulation of ACE2 expression. These alveolar macrophages act as the front line immune cells defending against the SARS-CoV-2 viral infection. This was supported by the ex vivo study with LPS-activated M1 macrophages. Recently, Ural et al. (2020) reported that alveolar macrophages displayed the phenotype of type 1 macrophages [49], while an interstitial subset of CD169^+^ lung-resident macrophages, primarily located around the airways in close proximity to the sympathetic nerves of the bronchovascular bundle, exhibit the characteristics of type 2 macrophages and anti-inflammatory effects [49]. Therefore, these two types of macrophages play an essential role in the immune surveillance and maintenance of homeostasis of the pulmonary system. Considering all current approaches for the prevention and treatment of COVID-19, there are no therapies, either being tested or at the beginning of the pipeline, that directly focus on the modulation of macrophages.

To address the overreaction of immune responses caused by viral infection, immune suppression regimens (e.g., chloroquine, hydroxychloroquine, JAK inhibitors, anti-cytokine therapy, and anti-IL-6R antibody) are being tested in clinical trials, which may be not sufficient to treat COVID-19 as they make subjects more vulnerable to the SARS-CoV-2 infection, in addition to their associated side effects. Rather, the immune modulation strategy like that of traditional Chinese herbs Lian-Hua-Qing-Wen Granule [50] may potentially enhance anti-viral immunity and improve clinical outcomes in patients after being infected with SARS-CoV-2. Additionally, over the last 10 years, Dr. Yong Zhao at Tianhe Stem Cell Biotechnologies has developed the Stem Cell Educator (SCE) therapy by utilizing the immune modulation of human cord blood-derived multipotent stem cells (CB-SC) for the treatment of type 1 diabetes and other inflammation-associated diseases. Clinical safety and efficacy have been demonstrated for SCE therapy through multicenter clinical trials [51–54]. Recently, mechanistic studies established that CB-SC-released exosomes could differentiate the purified CD14^+^ monocytes into M2 macrophages [15], suggesting that SCE therapy may have the potential to treat COVID-19 (ClinicalTrials.gov: NCT04299152). Targeting alveolar and other tissue macrophages through immune modulations may be potentially beneficial to correct the viral inflammation, effectively ameliorate anti-viral immunity, efficiently reduce the viral load, improve clinical outcomes, expedite the patient recovery, and decline the rate of mortality in patients after being infected with SARS-CoV-2.

## Acknowledgments

Authors are grateful to Mr. Poddar and Mr. Ludwig for generous funding support via Hackensack UMC Foundation.

## Authorship

Y.Z.: supervised experiments, and contributed to concepts, experimental design, data analysis and interpretation, manuscript writing, and final approval of manuscript; X.S.: performed most experiments and data analysis; W.H., performed the macrophage-associated experiments; H.Y.: performed flow cytometry on human primary epithelial cells; Ye. Z.: graphic design of Figure 6; L. Z.: manuscript writing and editing.

